# A Process Model Account of the Role of Dopamine in Intertemporal Choice

**DOI:** 10.1101/2022.10.24.513522

**Authors:** Alexander Soutschek, Philippe N. Tobler

## Abstract

Theoretical accounts disagree on the role of dopamine in intertemporal choice and assume that dopamine either promotes delay of gratification by increasing the preference for larger rewards or that dopamine reduces patience by enhancing the sensitivity to waiting costs. Here, we reconcile these conflicting accounts by providing empirical support for a novel process model according to which dopamine contributes to two dissociable components of the decision process, evidence accumulation and starting bias. We re-analyzed a previously published data set where intertemporal decisions were made either under the D2 antagonist amisulpride or under placebo by fitting a hierarchical drift diffusion model that distinguishes between dopaminergic effects on the speed of evidence accumulation and the starting point of the accumulation process. Blocking dopaminergic neurotransmission not only strengthened the sensitivity to whether a reward is perceived as worth the delay costs during evidence accumulation (drift rate) but also attenuated the impact of waiting costs on the starting point of the evidence accumulation process (bias). Taken together, our findings support a novel, process-based account of the role of dopamine for cost-benefit decision making, highlight the potential benefits of process-informed analyses and advance our understanding of dopaminergic contributions to decision making.

## Introduction

Many decisions require trading-off potential benefits (rewards) against the costs of actions, such as the time one has to wait for reward to occur (Soutschek & Tobler, 2018). The neurotransmitter dopamine is thought to play a central role in such cost-benefit trade-offs by increasing the tolerance for action costs in order to maximize net subjective benefits (Beeler, 2012; Robbins & Everitt, 1992; Salamone & Correa, 2012; Schultz, 2015). Tonic dopaminergic activity was hypothesized to implement a “cost control” which moderates whether a reward or goal is considered to be worth its costs (Beeler & Mourra, 2018). Prominent accounts of dopaminergic functioning thus predict that dopamine should strengthen the preference for costly larger-later (LL) over less costly smaller-sooner (SS) rewards. However, empirical studies modulating dopaminergic neurotransmission during intertemporal decision making provided inconsistent evidence for these hypotheses (for a review, see Webber, Lopez-Gamundi, Stamatovich, de Wit, & Wardle, 2020). Blocking dopaminergic activation even seems to increase rather than to reduce the preference for delayed outcomes (Arrondo et al., 2015; Soutschek et al., 2017; Wagner, Clos, Sommer, & Peters, 2020; Weber et al., 2016), in apparent contrast to accounts proposing that lower dopaminergic activity should decrease the attractiveness of costly rewards (Beeler & Mourra, 2018; Robbins & Everitt, 1992; Salamone & Correa, 2012). Thus, the link between dopamine and cost-benefit weighting in intertemporal choice remains elusive. Yet, a plausible account of how dopamine affects cost-benefit weighting is important given that deficits in delay of gratification belong to the core symptoms of several psychiatric disorders and that dopaminergic medication plays a central role in the treatment of these and other disorders (Hasler, 2012; MacKillop et al., 2011).

To account for the conflicting findings on the role of dopamine in intertemporal choice, recent proximity accounts hypothesized that dopamine – in addition to strengthening the pursuit of valuable goals – increases also the preference for proximate over distant rewards (first formulated by (Westbrook & Frank, 2018); see also (Soutschek, Jetter, & Tobler, 2022)). While proximity and action costs often correlate negatively (as cost-free immediate rewards are typically more proximate than costly delayed rewards), they can conceptually be distinguished: perceived costs depend on an individual’s internal state (e.g., available resources to wait for future rewards), whereas proximity is determined by situational factors like familiarity or concreteness (Westbrook & Frank, 2018). The hypothesis that dopamine increases the proximity advantage of sooner over later rewards is consistent with the observed stronger preference for LL options after D2R blockade, which could not be explained by standard accounts of the role of dopamine in cost-benefit decisions (Beeler & Mourra, 2018; Salamone & Correa, 2012).

Still, the question remains as to how the proximity account can be reconciled with the large body of evidence for a motivating role of dopamine in other domains than intertemporal choice (Webber et al., 2020). We recently suggested that both accounts may be unified within the framework of computational process models like the drift diffusion model (DDM) (Soutschek et al., 2022). DDMs assume that decision makers accumulate evidence for two reward options until a decision boundary is reached. The dopamine-mediated cost control may be implemented via dopaminergic effects on the evaluation of reward magnitudes and delay costs during the evidence accumulation process (drift rate), while a proximity advantage for sooner over delayed rewards may shift the starting bias towards the decision boundary for sooner rewards (Soutschek et al., 2022; Westbrook & Frank, 2018). Such proximity effects on the starting bias could reflect an automatic bias towards immediate rewards as posited by dual process models of intertemporal choice (Figner et al., 2010; McClure, Laibson, Loewenstein, & Cohen, 2004), whereas the influence of reward and delay on the drift rate involves more controlled and attention-demanding weighting of costs and benefits. Combining these two, in their consequences on overt choices partially opposing, but independent, effects of dopamine in a unified and tractable account could reconcile conflicting findings. In turn, such a process account might provide a knowledge basis to advance our understanding of the neurochemical basis of the decision making deficits in clinical disorders and improve the effectiveness of pharmaceutical interventions.

Here, we tested central assumptions of the proposed account by re-analyzing the data from a previous study that investigated how the dopamine D2 receptor antagonist amisulpride impacts cost-benefit weighting in intertemporal choice. Previously reported analyses of these data had shown no influence of D2R blockade on the mean preferences for LL over SS options (Soutschek et al., 2017). However, they had not asked whether the D2 antagonist moderates the influences of reward magnitudes and delay costs on subcomponents of the decision process within the framework of a drift diffusion model. We re-analyzed the data set with hierarchical Bayesian drift diffusion modelling to test two central assumptions of the proposed account on dopamine’s role in cost-benefit weighting. First, if D2R activation implements a cost threshold moderating the evaluation of whether a reward is worth the action costs, then blocking D2R activation with amisulpride should increase the influence of reward magnitude on the speed of evidence accumulation, with costly small rewards becoming less acceptable than under placebo. Second, if dopamine also moderates the impact of proximity on choices (which affects the starting bias rather than the speed of the evidence accumulation process), D2R blockade should attenuate the effects of waiting costs on the starting bias.

## Results

To disentangle how dopamine contributes to distinct subcomponents of the choice process, we re-analyzed a previously published data set where 56 participants had performed an intertemporal choice task under the D2 antagonist amisulpride (400 mg) and placebo in two separate sessions (Soutschek et al., 2017) (Figure 1). First, we assessed amisulpride effects on intertemporal choices with conventional model-based and model-free analyses, as they are employed by other pharmacological studies on cost-benefit weighting. Hyperbolic discounting of future rewards was not significantly different under amisulpride (mean log-k = −2.07) compared with placebo (mean log-k = −2.19), Bayesian t-test, HDI_mean_ = 0.21, HDI_95%_ = [−0.28; 0.70], and there were also no drug effects on choice consistency (inverse temperature), HDI_mean_ = −0.28, HDI_95%_ = [−0.71; 0.13]. Model-free Bayesian mixed generalized linear models (MGLMs) revealed a stronger preference for LL over SS options with increasing differences in reward magnitudes, HDI_mean_ = 6.32, HDI_95%_ = [5.03; 7.83], and with decreasing differences in delay of reward delivery, HDI_mean_ = −1.27, HDI_95%_ = [−1.87; −0.60]. The impact of delays on choices was significantly reduced under amisulpride compared with placebo, HDI_mean_ = 0.75, HDI_95%_ = [0.02; 1.67] (Table 1). Thus, based on these conventional analyses one would conclude that reduction of D2R neurotransmission lowers the sensitivity to delay costs, which on the one hand agrees with one line of previous findings (Arrondo et al., 2015; Wagner et al., 2020; Weber et al., 2016). On the other hand, this result seems to contradict the widely held assumption that dopamine increases the preference for costly over cost-free outcomes (Beeler & Mourra, 2018; Webber et al., 2020; Westbrook et al., 2020), because according to this view lower dopaminergic activity should increase, rather than decrease, the impact of waiting costs on LL choices. However, analyses that consider only the observed choices do not allow disentangling dopaminergic influences on distinct subcomponents of the choice process.

**Table 1.**
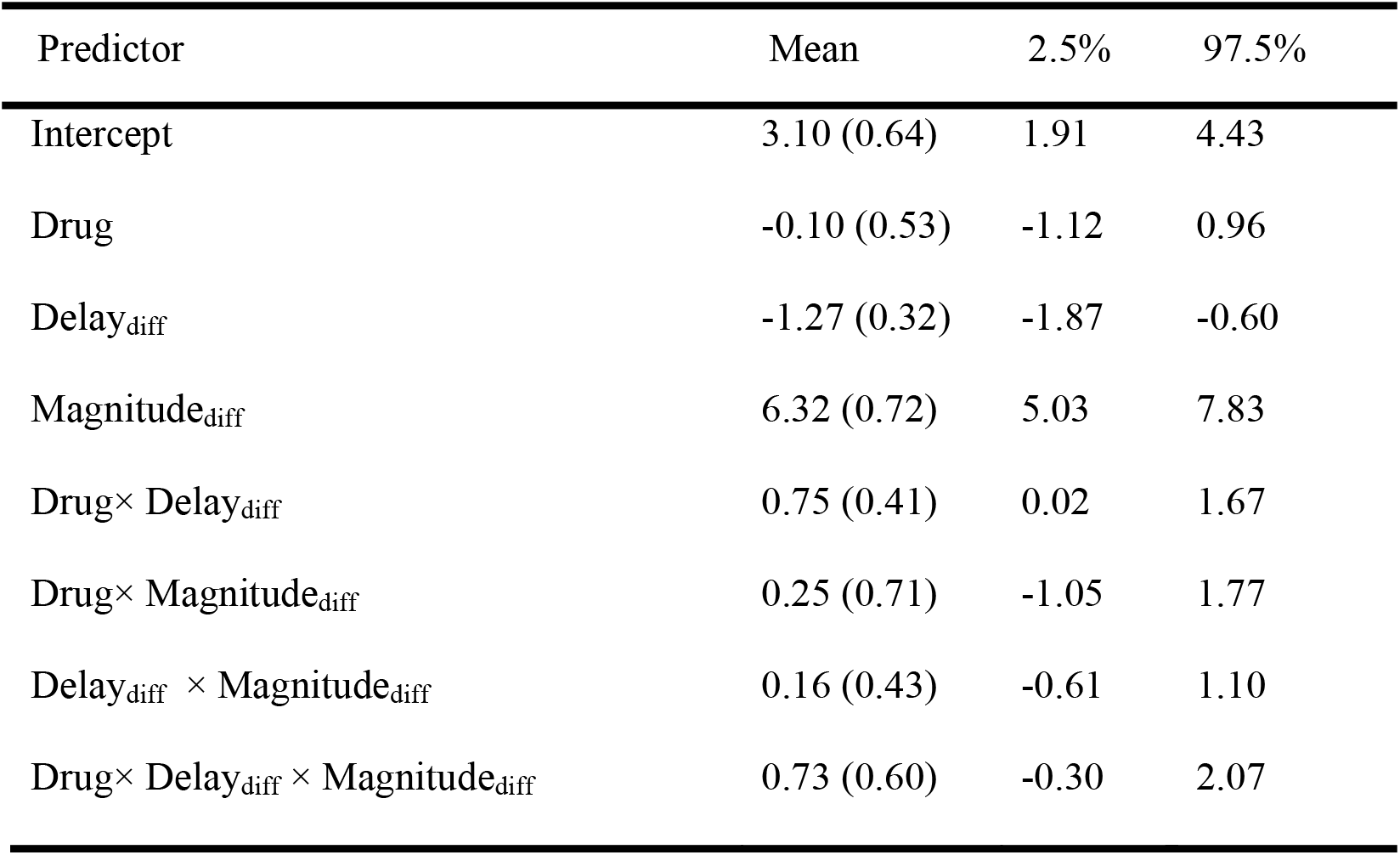
Results of Bayesian generalized linear model regressing binary choices (LL versus SS option) on predictors for Drug, Magnitude_diff_, Delay_diff_, and the interaction terms. Standard errors of the mean of the posterior distributions are in brackets.

**Figure 1.**
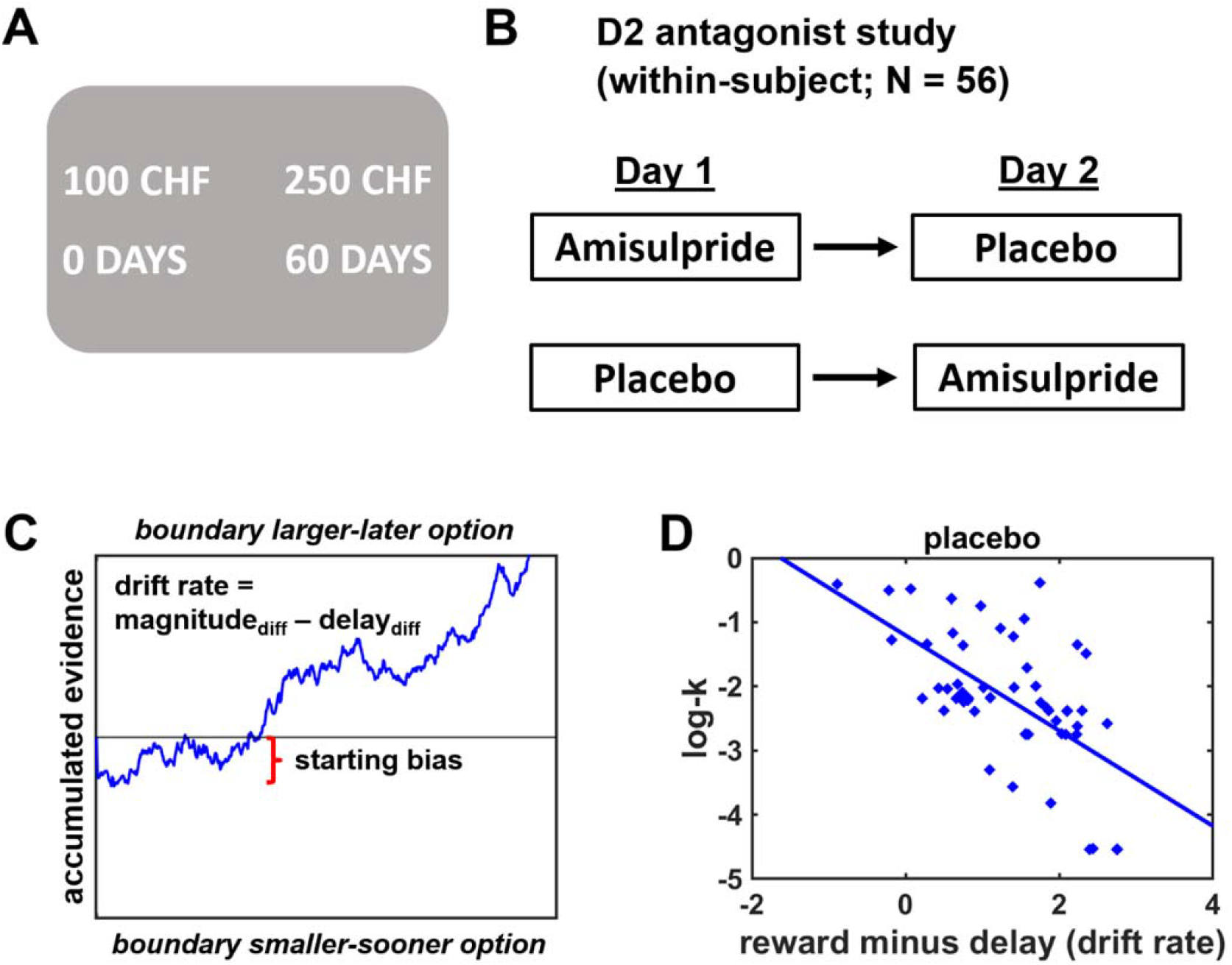
Task design and experimental procedures. (A) Participants made choices between alternatives that provided smaller sooner rewards (e.g., 100 Swiss francs in 0 days) or larger later rewards (e.g., 250 Swiss francs in 60 days). (B) In a double-blind crossover design, participants performed the intertemporal decision task after administration of the D2 antagonist amisulpride or placebo on two separate days. (C) Illustration of the choice process in the framework of a drift-diffusion model. After a non-decision time τ (not shown here), evidence is accumulated from a starting point ζ with the weighted difference between benefits and action costs determining the speed of the accumulation process (drift rate *v*) towards the boundaries for the larger-later or smaller-sooner option. (D) Individual differences in delay discounting under placebo (log-k) decrease with the difference in weights assigned to rewards and delay costs during evidence accumulation, replicating previous findings (Amasino, Sullivan, Kranton, & Huettel, 2019).

DDMs paint a fuller picture of the decision process than pure choice data by integrating information from observed choices and decision times. DDMs assume that agents accumulate evidence for the choice options (captured by the drift parameter *v*) from a starting point ζ until the accumulated evidence reaches a decision threshold (boundary parameter *a*; Figure 1C). Following previous procedures analyzing intertemporal choices with drift diffusion models (Amasino et al., 2019), we assumed that the drift rate ν integrates reward magnitudes and delays of choice options via attribute-wise comparisons (DDM-1). In addition, we allowed also the starting bias to vary as a function of differences in delay costs, in line with recent proximity accounts of dopamine (Westbrook & Frank, 2018).

A sanity check revealed that larger differences between the reward magnitudes of the LL and SS options bias evidence accumulation towards the LL option, HDI_mean_ = 2.41, HDI_95%_ = [1.93; 2.95], whereas larger differences in delays bias accumulation in favor of the SS option, HDI_mean_ = −1.13, HDI_95%_ = [−1.53; −0.78]. Moreover, we assessed the relationship between the difference in DDM parameters (reward magnitude – delay) and hyperbolic discount parameters log-k as purely choice-based indicator of impulsiveness. Replicating previous findings, we found that the differences in the weights relate to individual differences in delay discounting, r = −0.61, *p* < 0.001 (Amasino et al., 2019) (Figure 1D), such that individuals weighting reward magnitudes more strongly over delays make more patient choices. Thus, our model parameters capture essential subprocesses of intertemporal decision making.

Next, we tested the impact of our dopaminergic manipulation on evidence accumulation: D2R blockade strengthened the impact of differences in reward magnitude on evidence accumulation, Drug × Magnitude_diff_: HDI_mean_ = 0.81, HDI_95%_ = [0.04; 1.71], while the contribution of differences in delay costs remained unchanged, Drug × Delay_diff_: HDI_mean_ = −0.30, HDI_95%_ = [−0.85; 0.20] (Figure 2A, B and Table 2). The drug-induced increase in sensitivity to variation in reward magnitude suggests that low rewards are considered less valuable under amisulpride compared with placebo (Figure 2C). This finding is consistent with the cost control hypothesis (Beeler & Mourra, 2018) according to which low dopamine levels reduce the attractiveness of smaller, below-average rewards.

**Table 2.** Results of hierarchical DDM-1. SEM: Standard errors of the mean of the posterior distributions.

| Parameter | Regressor | Mean (SEM) | 2.5% | 97.5% |
| --- | --- | --- | --- | --- |
| Drift rate | Delay<sub>diff</sub> | -1.13 (0.19) | -1.53 | -0.78 |
|  | Magnitude<sub>diff</sub> | 2.41 (0.26) | 1.93 | 2.95 |
|  | Drug×Delay<sub>diff</sub> | -0.30 (0.27) | -0.85 | 0.20 |
|  | Drug×Magnitude<sub>diff</sub> | 0.81 (0.42) | 0.04 | 1.71 |
|  | V<sub>max</sub> | 0.80 (0.05) | 0.70 | 0.90 |
|  | Drug×V<sub>max</sub> | -0.00 (0.06) | -0.12 | 0.12 |
| Decision threshold | Placebo | 3.96 (0.15) | 3.67 | 4.26 |
|  | Drug | 0.17 (0.15) | -0.11 | 0.46 |
| Starting bias | Placebo | 0.57 (0.01) | 0.54 | 0.60 |
|  | Drug | -0.04 (0.02) | -0.08 | -0.001 |
|  | Delay<sub>diff</sub> | -0.02 (0.01) | -0.03 | -0.002 |
|  | Drug×Delay<sub>diff</sub> | 0.02 (0.01) | 0.001 | 0.04 |
| Non-decision time | Placebo | 1.48 (0.06) | 1.36 | 1.60 |
|  | Drug | -0.10 (0.07) | -0.24 | 0.03 |

**Figure 2.**
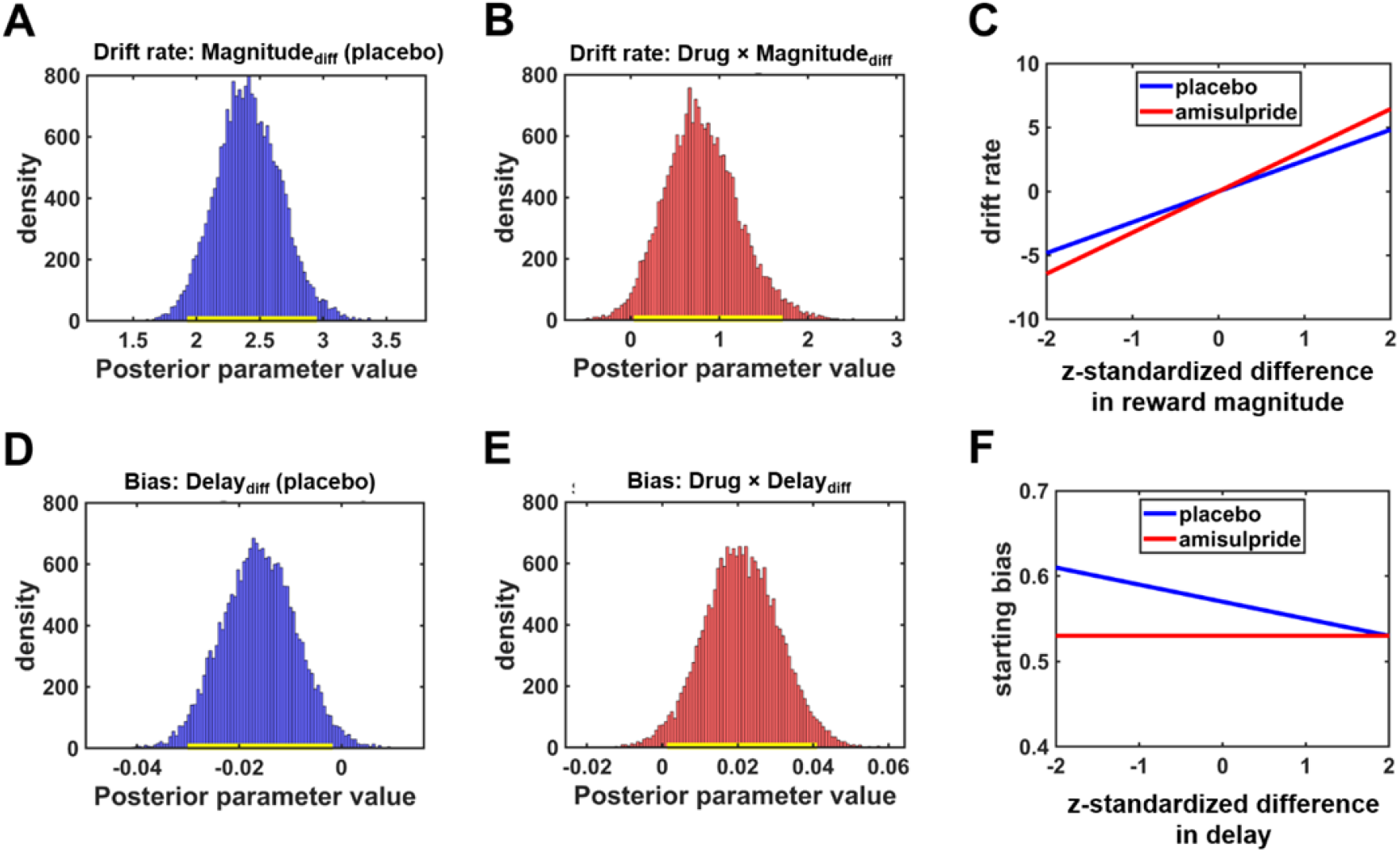
D2R blockade affects multiple components of the intertemporal decision process. (A) Larger differences in reward magnitude between the LL and SS option increased drift rates, speeding up evidence accumulation towards LL options under placebo. (B) The impact of differences in reward magnitude was significantly stronger under amisulpride than under placebo. (C) Drug-dependent impact of differences in reward magnitude on the drift rate. Because the sensitivity to differences in reward magnitude was stronger under amisulpride than under placebo (steeper slope), D2R blockade speeded-up evidence accumulation towards the boundary for LL choices if differences in reward magnitude between the LL and SS options were large. In contrast, if the difference in reward magnitude was small, the drift rate was more negative under amisulpride compared with placebo, speeding up evidence accumulation towards the SS option. (D) The starting point of the evidence accumulation process was increasingly shifted towards the SS option for larger differences in waiting costs under placebo. (E) This impact of delay costs on the starting bias was significantly reduced under amisulpride. As illustrated in (F), reducing dopaminergic action on D2R with amisulpride shifted the starting bias towards the boundary for the SS option predominantly if no option possessed a clear proximity advantage (small difference between delays). In A, B, D, and E, yellow bars close to x-axis indicate 95% HDIs.

When we assessed dopaminergic effects on the starting bias, we observed that under placebo increasing differences in delay shifted the starting point towards the SS option, HDI_mean_ = 0.81, HDI_95%_ = [0.04; 1.71], suggesting that the bias parameter is closer to the proximate (SS) option the stronger the proximity advantage of the SS over the LL option. Amisulpride shifted the starting bias towards the SS option for smaller differences in delay, main effect of Drug_y_: HDI_mean_ = −0.04, HDI_95%_ = [−0.08; 0.01], but also attenuated the impact of delay, Drug × Delay_diff_: HDI_mean_ = 0.02, HDI_95%_ = [0.001; 0.04]. Thus, dopamine appears to moderate the impact of temporal proximity on the starting bias (Figure 2D-F), providing support for recent proximity accounts of dopamine (Soutschek et al., 2022; Westbrook & Frank, 2018). Moreover, compared to the model-free analysis, our process model (which uses not only binary choice but also response time data) provides a fuller picture of the subcomponents of the choice process affected by the dopaminergic manipulation.

Next, we investigated the relation between the drug effects on the drift rate and on the starting bias. We found no evidence that the two effects correlated, *r* = 0.07, *p* = 0.60, suggesting that amisulpride effects on these subprocesses were largely independent of each other. Control analyses revealed no effects of amisulpride on non-decision times, HDI_mean_ = −0.10, HDI_95%_ = [−0.24; 0.03], or the decision threshold, HDI_mean_ = 0.17, HDI_95%_ = [−0.11; 0.46]. Thus, the results of DDM-1 suggest that dopamine moderates the influence of choice attributes on both the speed of evidence accumulation and on the starting bias, consistent with recent accounts (Soutschek et al., 2022; Westbrook & Frank, 2018) of dopamine’s role in cost-benefit weighting.

To test the robustness of our DDM findings, we computed further DDMs where we either removed the impact of Delay_diff_ on the starting bias (DDM-2) or the impact of Magnitude_diff_ and Delay_diff_ on the drift rate (DDM-3). Importantly, our original DDM-1 (DIC = 9,478) explained the data better than DDM-2 (DIC = 9,481) or DDM-3 (DIC = 10,224). Nevertheless, amisulpride moderated the impact of Magnitude_diff_ on the drift rate also in DDM-2, HDI_mean_ = 0.86, HDI_95%_ = [0.18; 1.64], and lowered the impact of Delay_diff_ on the starting bias in DDM-3, HDI_mean_ = −0.02, HDI_95%_ = [−0.04; −0.001]. Thus, the dopaminergic effects on these subcomponents of the choice process are robust to the exact specification of the DDM.

We compared the winning account also with alternative process models of intertemporal choice. While in DDM-1 the drift rate depends on separate comparisons between choice attributes, one might alternatively assume that they compare the discounted subjective reward values of both options (Wagner et al., 2020), as given by the hyperbolic discount functions. However, a DDM where the drift rate was modelled as the difference between the hyperbolically discounted reward values (with the discount factor as free parameter; DDM-4) showed a worse model fit (DIC = 10,720) than DDM-1. This replicates previous findings according to which intertemporal choices can better be explained by attribute-wise than by option-wise comparison strategies (Amasino et al., 2019; Dai & Busemeyer, 2014; Reeck, Wall, & Johnson, 2017).

Next, we investigated an alternative to the proposal that differences in delay affect the starting bias via proximity effects. Specifically, we tested whether evidence for delay costs are accumulated earlier than for reward magnitude (relative-starting-time (rs)DDM (Amasino et al., 2019; Lombardi & Hare, 2021)). From the perspective of rsDDMs, evidence accumulation for delays would start after a shorter non-decision time than for rewards, which is expressed by the variable τ_diff_ (if τ_diff_ > 0, non-decision time is shorter for delays than rewards, and vice versa if τ_diff_ < 0). However, also this rsDDM (DDM-5) explained the data less well (DIC = 9,548) than DDM-1. Thus, DDM-1 explains the current data better than alternative DDMs.

The currently used dose of amisulpride (400 mg) is thought to have predominantly postsynaptic effects on D2Rs, while lower doses (50-300 mg) might show presynaptic rather than postsynaptic effects (Schoemaker et al., 1997). Given that we used the same dose in all participants, one might argue that we may have studied presynaptic effects in individuals with relatively high body mass (which lowers the effective dose). However, we observed no evidence that individual random coefficients for the drug effects on the drift rate or on the starting bias correlated with body weight, all r < 0.15, all *p* > 0.28. There was thus no evidence that pharmacological effects on intertemporal choices depended on body weight as proxy of effective dose, which provides no support for the possibility that the reported results reflect pre-synaptic effects.

As further check of the explanatory adequacy of DDM-1, we performed posterior predictive checks and parameter recovery analyses. Plotting the observed RTs (split into quintiles according to Magnitude_diff_ and Delay_diff_) against the simulated RTs based on the parameter estimates from the different DDMs suggests that the DDMs provide reasonable accounts of the observed data (Figure 3). Moreover, the squared differences between observed and simulated RTs were smaller for DDM-1 (0.83) than for alternative DDMs (DDM-2: 0.85; DDM-3: 0.98; DDM-4: 0.89; DDM-5: 1.63). To assess parameter recovery, we re-computed DDM-1 on ten simulated data sets based on the original DDM-1 parameter estimates. All group-level parameters from the simulated data were within the 95% HDI of the original parameter estimates, except for the non-decision time tau (which suggests that our model tends to overestimate the duration of decision-unrelated processes). Nevertheless, all parameters determining the outcome of the decision process (i.e., the choice made) as well as the dopaminergic effects on the parameters could reliably be recovered by DDM-1.

**Figure 3.**
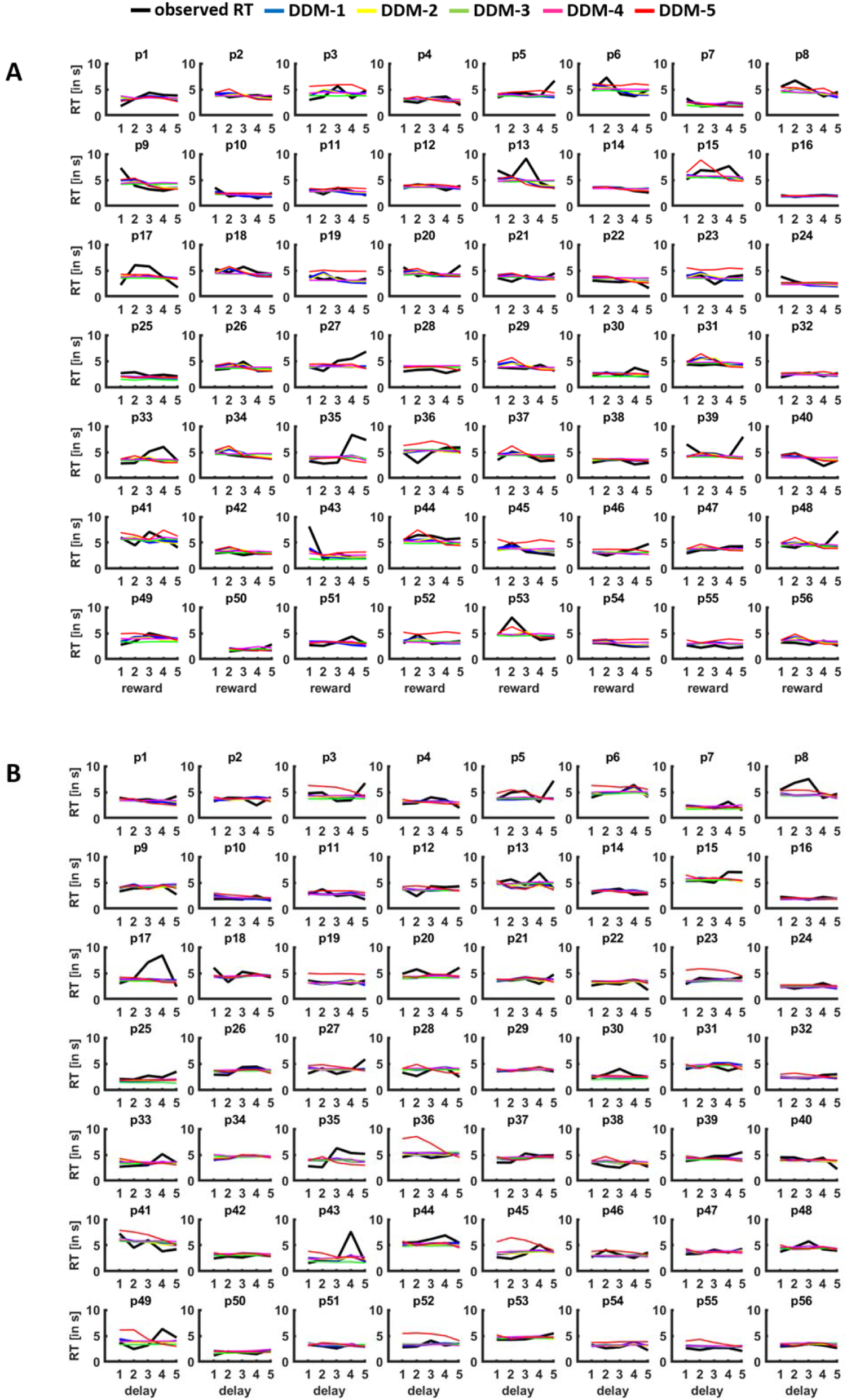
Posterior predictive checks. For each individual participant (p1-p56), observed RTs (in black) are plotted against the RTs simulated based on the parameters for DDM 1-5, separately for differences in (A) reward magnitude and (B) delay (quintiles). The plots suggest that the DDMs provide reasonable accounts of the observed RTs.

## Discussion

Dopamine is hypothesized to play a central role in human cost-benefit decision making, but existing empirical evidence does not conclusively support the widely shared assumption that dopamine promotes the pursuit of high benefit-high cost options (for reviews, see (Soutschek et al., 2022; Webber et al., 2020)). By manipulating dopaminergic activity with the D2 antagonist amisulpride, we provide empirical evidence for a novel process model of cost-benefit weighting that reconciles conflicting views by assuming dissociable effects of dopamine on distinct subcomponents of the decision process.

D2R blockade (relative to placebo) increased the sensitivity to variation in reward magnitudes during evidence accumulation, such that only relatively large future rewards were considered to be worth the waiting cost, whereas small delayed rewards were perceived as less valuable than sooner rewards. This dopaminergic impact on the drift rate is consistent with the view that D2R-mediated tonic dopamine levels implement a cost control determining whether a reward is worth the required action costs (Beeler & Mourra, 2018). From this perspective, lowering D2R activity with amisulpride resulted in a stricter cost control such that only rather large delayed rewards were able to overcome D2R-mediated cortical inhibition (Lerner & Kreitzer, 2011). While this effect is consistent with the standard view according to which dopamine increases the preference for large costly rewards (Robbins & Everitt, 1992; Salamone & Correa, 2012; Schultz, 2015), the dopaminergic effects on the starting bias parameter yielded a different pattern. Here, inhibition of D2R activation reduced the impact of delay costs on the starting bias, such that for shorter delays (where the immediate reward has only a small proximity advantage) D2R inhibition shifts the bias towards the SS option. This finding represents first evidence for the hypothesis that dopamine moderates the impact of proximity (e.g., more concrete versus more abstract rewards) on cost-benefit decision making (Soutschek et al., 2022; Westbrook & Frank, 2018).

Conceptually, the assumption of proximity effects on the starting bias is consistent with dual process models of intertemporal choice assuming that individuals are (at least partially) biased towards selecting immediate over delayed rewards (Figner et al., 2010; McClure et al., 2004). This automatic favoring of immediate rewards is reflected in a shift of the starting bias and thus occurs before the evidence accumulation process, which relies on attention-demanding cost-benefit weighting (Zhao, Diederich, Trueblood, & Bhatia, 2019). In agreement with this notion, we DDM-1 with temporal proximity-dependent bias showed better fit than DDM-5 with variable non-decision times for rewards and delays.

A dopaminergic modulation of proximity effects provides an elegant explanation for the fact that in most D2 antagonist studies D2R reduction increased the preference for LL options (Arrondo et al., 2015; Soutschek et al., 2017; Wagner et al., 2020; Weber et al., 2016), contrary to the predictions of energization accounts (Beeler & Mourra, 2018; Salamone & Correa, 2012). Noteworthy, the dopaminergic effects on evidence accumulation and on the starting bias promote potentially different action tendencies, as the impact of amisulpride on evidence accumulation lowered the weight assigned to small future rewards, whereas the amisulpride effects on the starting bias increased the likelihood of LL options being chosen. Rather than generally biasing impulsive or patient choices, the impact of dopamine on decision making may therefore crucially depend on the rewards at stake and the associated waiting costs (Figure 4). In our model, lower dopamine levels strengthen the preference for high reward-high cost options predominantly two situations. First, if differences in reward magnitude are high (e.g., choosing between your favorite meal versus a clearly less liked dish) and, second, if the less costly option has a clear proximity advantage over the costlier one (having dinner in a restaurant close-by or a preferred restaurant on the other side of town). Conversely, if differences in both expected reward and waiting costs are small, lower dopamine may bias choices in favor of low-cost rewards over high-cost rewards. By extension, higher dopamine levels should increase the preference for an SS option if the SS option has a pronounced proximity advantage over the LL option, and bias the acceptance of LL options if both options are associated with similar waiting costs. We note though that the effects of increasing dopamine levels are less predictable than the effects of lowering dopaminergic activity due to possible inverted-u shaped dopamine-response curves (Floresco, 2013); potentially, the dopaminergic effects on drift rate and starting bias might even follow different dose-response functions. Taken together, our process model of the dopaminergic involvement in cost-benefit decisions allows reconciling conflicting theoretical accounts and (apparently) inconsistent empirical findings by showing that dopamine moderates the effects of reward magnitudes and delay costs on different subcomponents of the choice process.

**Figure 4.**
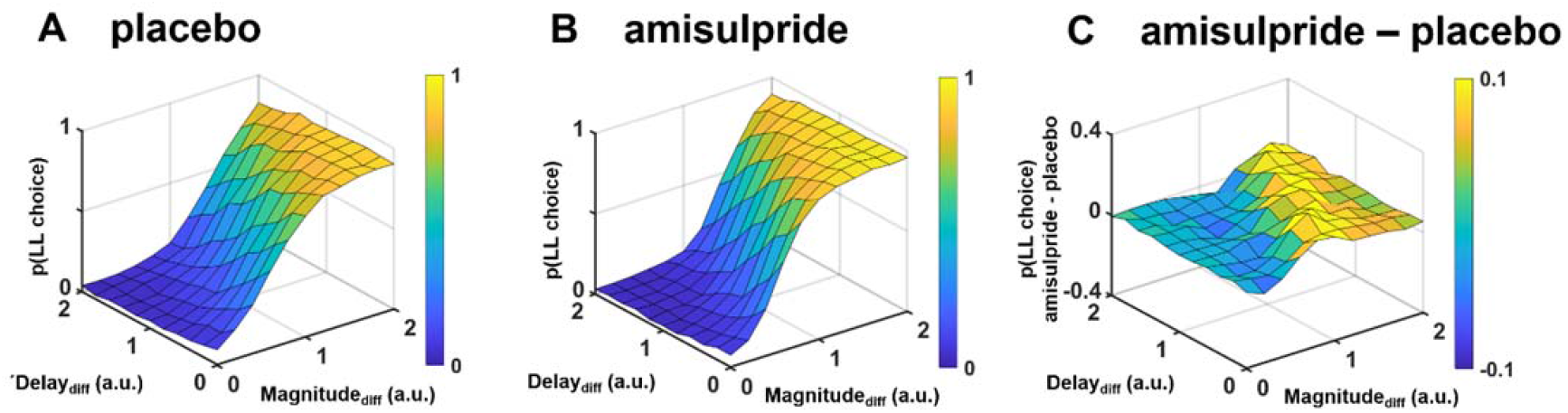
Illustration of how dopaminergic effects on intertemporal choices depend on differences in both reward magnitude and delay in the proposed framework, separately for (A) placebo, (B) amisulpride, and (C) the difference between amisulpride and placebo. Plots are based on simulations assuming the group-level parameter estimates we observed under placebo and amisulpride. As dopaminergic effects on decision making affect both reward processing (via the drift rate) and cost processing (via the starting bias), the specific combination of rewards and delays determines whether D2R blockade increases or decreases the probability of LL choices. Low dopamine levels reduce the proximity advantage of SS over LL options particularly if differences in action costs between reward options are large, promoting choices of the LL option. In contrast, if no option possesses a proximity advantage (small differences between delays), dopaminergic effects on evidence accumulation dominate, such that the LL option is perceived as less worth the waiting costs, particularly if its reward magnitude differs only little from that of the alternative SS option.

We note that the moderating roles of differences in delays are also reflected in the significant interaction between drug and delay from the model-free analysis, although this analysis could provide no insights into which subcomponents of the choice process are affected by dopamine. As the influence of dopamine on decision making varies as a function of the differences in reward magnitude and waiting costs, the outcomes of standard analyses like mean percentage of LL choices or hyperbolic discount parameters may be specific to the reward magnitudes and delays administered in a given study. For example, if an experimental task includes large differences between rewards and delays, dopamine antagonists may reduce delay discounting, whereas studies with smaller differences between these choice attributes may observe no effect of dopaminergic manipulations (Figure 4). Standard analyses that measure patience by one behavioral parameter only (e.g., discount factors) may thus result in misleading findings. In contrast, process models of decision making do not just assess whether a neural manipulation increases or reduces patience; instead, they quantify the influence of a manipulation on the weights assigned to rewards and waiting costs during different phases of the choice process, with these weights being less sensitive to the administered choice options in a given experiment. Process models may thus provide a less option-specific picture of the impact of pharmacological and neural manipulations.

As potential alternative explanation for the enhanced influence of reward magnitude under amisulpride, one might argue that D2R blockade generally increases the signal-to-noise ratio for decision-relevant information. However, this notion is inconsistent with the proposed role of D2R activation for precise action selection (Keeler, Pretsell, & Robbins, 2014), because this view would have predicted amisulpride to result in noisier (less precise action selection) rather than less noisy evidence accumulation. Moreover, our data provide no evidence for drug effects on the inverse temperature parameter measuring choice consistency, and there were also no significant correlations between amisulpride effects on reward and delay processing, contrary to what one should expect if these effects were driven by the same mechanism. We also note that while higher doses of amisulpride (as administered in the current study) antagonize post-synaptic D2Rs, lower doses (50-300 mg) were found to block action at pre-synaptic dopamine receptors (Schoemaker et al., 1997), which may result in amplified phasic dopamine release and thus increased sensitivity to benefits (Frank & O’Reilly, 2006). If such pre-synaptic mechanisms explained the observed amisulpride effects on evidence accumulation, the strength of these effects should increase with body weight (as higher body weight lowers the effective dose). However, we observed no significant correlations between individual drug effects and body weight. This appears at variance with the assumption that the observed drug effects might result from presynaptic versus postsynaptic mechanisms of action.

To conclude, our findings may shed a new light on the role of dopamine in psychiatric disorders that are characterized by deficits in impulsiveness or cost-benefit weighting in general (Hasler, 2012), and where dopaminergic drugs belong to the standard treatments for deficits in value-related and other behavior. Dopaminergic manipulations yielded mixed results on impulsiveness in psychiatric and neurologic disorders (Acheson & de Wit, 2008; Antonelli et al., 2014; Foerde et al., 2016; Kayser, Vega, Weinstein, Peters, & Mitchell, 2017), and our process model regarding the role of dopamine for delaying gratification explains some of the inconsistencies between empirical findings (on top of factors like non-linear dose-response relationships). As similarly inconsistent findings were observed also in the domains of risky and social decision making (Soutschek et al., 2022; Webber et al., 2020), the proposed process model may account for the function of dopamine in these domains of cost-benefit weighting as well. By deepening the understanding of the role of dopamine in decision making, our findings provide insights into how abnormal dopaminergic activation, and its pharmacological treatment, in psychiatric disorders may affect distinct aspects of decision making.

## Materials and Methods

### Participants

In a double-blind, randomized, within-subject design, 56 volunteers (27 female, M_age_ = 23.2 years, SD_age_ = 3.1 years) received 400 mg amisulpride or placebo in two separate sessions (two weeks apart) as described previously (Soutschek et al., 2017). Participants gave informed written consent before participation. The study was approved by the Cantonal ethics committee Zurich.

### Task design

In both sessions, participants made intertemporal decisions 90 min after drug intake. We used a dynamic version of a delay discounting task in which the choice options were individually selected such that the information provided by each decision was optimized (dynamic experiments for estimating preferences, DEEP; (Toubia, Johnson, Evgeniou, & Delquie, 2013)). On each trial, participants decided between an SS (reward magnitude 5-250 Swiss francs, delay 0-30 days) and an LL option (reward magnitude 15-300 Swiss francs, delay 3-90 days). Participants pressed the left or right arrow keys on a standard keyboard to choose the option presented on the left or right side of the screen. On each trial, the reward options were presented until participants made a choice. The next choice options were displayed after an intertrial interval of 1 s. Participants made a total of 20 choices between SS and LL options.

### Statistical analysis

#### Drift-diffusion modelling

We analysed drug effects on intertemporal decision making with hierarchical Bayesian drift diffusion modelling using the JAGS software package (Plummer, 2003). JAGS utilizes Markov Chain Monte Carlo sampling for Bayesian estimation of drift diffusion parameters (drift rate ν, boundary α, bias ζ, and non-decision time τ) via the Wiener module (Wabersich & Vandekerckhove, 2014) on both the group- and the participant-level. In our models, the upper boundary (decision threshold) was associated with a choice of the LL option, the lower boundary with a choice of the SS option. A positive drift rate thus indicates evidence accumulation towards the LL option, a negative drift rate towards the SS option. As we were interested in how dopamine modulates different subcomponents of the choice process, in DDM-1 we assumed that the drift rate v is influenced by the comparisons of reward magnitudes and delays between the SS and LL options (Amasino et al., 2019; Dai & Busemeyer, 2014):

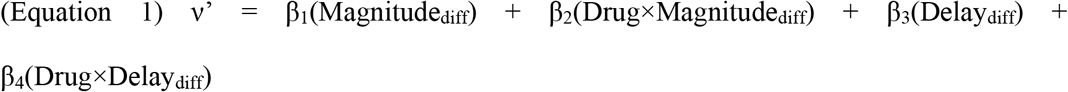

Magnitude_diff_ indicates the difference between the reward magnitudes of the LL and SS options, Delay_diff_ indicates the difference between the corresponding delays. Both Magnitude_diff_ and Delay_diff_ were z-transformed to render the size of the parameter estimates comparable (Amasino et al., 2019). Following previous procedures, we transformed v’ with a sigmoidal link function as this procedure explains observed behaviour better than linear link functions (Fontanesi, Gluth, Spektor, & Rieskamp, 2019; Wagner et al., 2020). Indeed, also the current data were better explained by a DDM with (DIC = 9,478) than without (DIC = 10,283) a sigmoidal link:

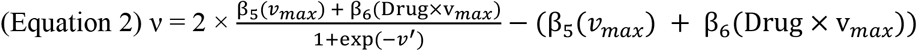

Next, we assessed whether delay costs affect the starting bias parameter ζ, as assumed by proximity accounts (Soutschek et al., 2022; Westbrook & Frank, 2018):

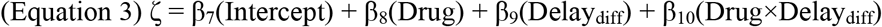

We also investigated whether the drug affected the decision threshold parameter α (equation 4) or the non-decision time τ (equation 5):

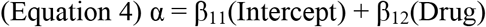

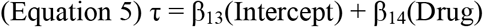

As the experiment followed a within-subject design, we modelled all parameters both on the group-level and on the individual level by assuming that individual parameter estimates are normally distributed around the mean group-level effect with a standard deviation λ (which was estimated separately for each group-level effect). We tested for significant effects by checking whether the 95% HDIs of the posterior samples of group-level estimates contained zero. Note that all statistical inferences were based on assessment of group-level estimates, as individual estimates might be less reliable due to the limited number of trials for each participant. We excluded the trials with the 2.5% fastest and 2.5% slowest response times to reduce the impact of outliers on parameter estimation (Amasino et al., 2019; Wagner et al., 2020). As priors, we assumed standard normal distributions for all group level effects (with mean = 0 and standard deviation = 1) and gamma distributions for λ (Wagner et al., 2020). For model estimation, we computed 2 chains with 500,000 samples (burning = 450,000, thinning = 5). 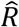 was used to assess model convergence in addition to visual inspection of chains. For all effects, 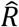 was below 1.01, indicating model convergence.

We compared DDM-1 also with alternative process models. DDM-2 was identical to DDM-1 but did not estimate starting bias as free parameter, assuming ζ = 0.5 instead, whereas DDM-3 left out the influences of Magnitude_diff_ and Delay_diff_ on the drift rate. In DDM-4 we assumed that the drift rate depends on the comparison of the hyperbolically discounted subjective values of the two choice options rather than on the comparison of choice attributes (Konovalov & Krajbich, 2019). In particular, the drift rate ν’ (prior to being passed through the sigmoidal link function) was calculated with:

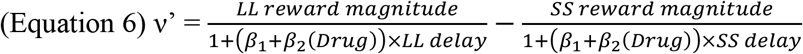

Here, β_1_ corresponds to the hyperbolic discount factor, which determines the hyperbolically discounted subjective values of the available choice options.

Finally, we considered a model without influence of Delay_diff_ on the starting bias but with separate non-decision times for rewards and delays. In more detail, DDM-5 included an additional parameter τ_diff_ which indicated whether the accumulation process started earlier for delays than for rewards (τ_diff_ > 0) or vice versa (τ_diff_ < 0). For example, if τ_diff_ > 0, evidence accumulation for delays starts directly after the non-decision time τ, whereas the accumulation process for reward magnitudes starts at τ + τ_diff_ (and then influences the driftrate together with Delay_diff_ until the decision boundary is reached). A recent study showed that such time-varying drift rates can be calculated as follows (Lombardi & Hare, 2021):

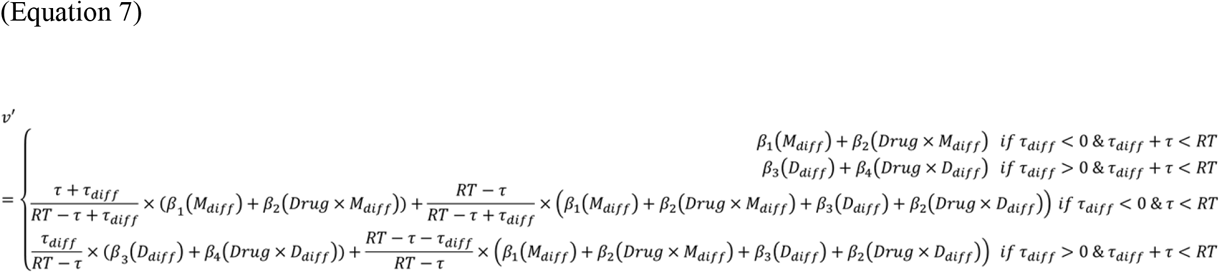

For the ease of reading, Magnitude_diff_ and Delay_diff_ are abbreviated as M_diff_ and D_diff_, respectively.

#### Posterior predictive checks and parameter recovery analyses

We performed posterior predictive checks to assess whether the DDMs explained key aspects of the empirical data. For this purpose, we simulated 1,000 RTs distributions based on the individual parameter estimates from all DDMs. We then binned trials into quintiles based on differences in reward magnitude and plotted the observed empirical data and the simulated data (averaged across the 1,000 simulations) as a function of these bins, separately for each individual participant. We performed the same analysis by binning trials based on differences in delay instead of reward magnitude.

We conducted a parameter recovery analysis by re-computing DDM-1 on ten randomly selected data sets which were simulated based on the original DDM-1 parameters. We checked parameter recovery by assessing whether group-level parameters from the simulated data lie within the 95% HDI of the original parameter estimates.

#### Model-free analyses

We analysed choice data also in a model-free manner and with a hyperbolic discounting model. In the model-free analysis, we regressed choices of LL versus SS options on fixed-effect predictors for Drug, Magnitude_diff_, Delay_diff_, and the interaction terms using Bayesian mixed models as implemented in the brms package in R (Bürkner, 2017). All predictors were also modelled as random slopes in addition to participant-specific random intercepts. Finally, the hyperbolic discounting model was fit using the hBayesDM toolbox (Ahn, Haines, & Zhang, 2017), using a standard hyperbolic discounting function:

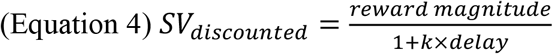

To translate subjective value into choices, we fitted a standard softmax function to each participant’s choices:

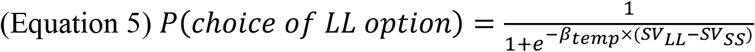

We estimated parameters capturing the strength of hyperbolic discounting (k) and choice consistency (*β*_*temp*_) separately for each participant and experimental session by computing two chains of 4,000 iterations (burning = 2,000). We then performed a Bayesian t-test on the log-transformed individual parameter estimates under placebo versus amisulpride using the BEST package (Kruschke, 2013).

## Acknowledgements

PNT received funding from the Swiss National Science Foundation (Grants 100019_176016, 100014_165884, and CRSII5_177277) and from the Velux Foundation (Grant 981). AS received an Emmy Noether fellowship (SO 1636/2-1) from the German Research Foundation.

## Conflict of interest statement

The authors declare to have no conflicts of interest.

## Data availability statement

The data supporting the findings of this study and the data analysis code will be available on Open Science Framework (https://osf.io/dp2me/; reviewer link: https://osf.io/dp2me/?view_only=f64654dd5d4942bb86cc2505863a8210).

## Notes

### Competing Interest Statement

The authors have declared no competing interest.

